# Metabolite profiles distinguish exposure to Dengue and Zika flaviviruses in human induced pluripotent stem cells (hiPSCs)

**DOI:** 10.64898/2026.01.29.702537

**Authors:** Tahira Fatima, Khyati Y. Mehta, Aaron Scholl, Bingjie Li, Djawed Bennouna, Maria Rios, Ewy A. Mathé, Sandip De

## Abstract

Flaviviruses such as Dengue virus (DENV), Zika virus (ZIKV) and West Nile Virus (WNV) pose growing public health threats, exacerbated by asymptomatic infections and non-vector transmission. Current diagnostics, including nucleic acid tests and serology, are limited by transient viremia and antibody cross-reactivity, underscoring the need for improved detection methods. We utilized untargeted metabolomics to differentiate DENV and ZIKV infections in human induced pluripotent stem cells (hiPSCs), a model relevant to study virus–host interactions, drug screening, and therapeutic safety. LC-MSbased profiling revealed virus-specific metabolic reprogramming during both acute and long-term infections. DENV3 induced early and sustained activation of metabolic features, while ZIKV-MR766 triggered transient suppression. Long term infection with DENV2, DENV3, or ZIKV-PRV resulted in distinct metabolic signatures in hiPSCs, with ZIKV-PRV showing the greatest divergence. Shared metabolite changes across conditions included amino acids (e.g., tryptophan, glutamate), lipids, and nucleosides. Functional studies demonstrated that tryptophan metabolism regulates infection dynamics: inhibiting serotonin biosynthesis reduced viral load, whereas blocking kynurenine synthesis enhanced viral replication. These findings position metabolomics as a viable approach for flavivirus detection and highlight metabolic pathways as potential therapeutic targets.

**Importance:** Accurate diagnosis of flavivirus infections remains challenging due to short periods of viremia, high rates of asymptomatic infection, and extensive serologic cross-reactivity among related viruses such as DENV and ZIKV. These limitations complicate clinical diagnosis, surveillance, and screening of human donor tissues. In this study, we demonstrate that untargeted metabolomics can distinguish ZIKV and DENV infections by identifying virus-specific host metabolic signatures during both acute and long-term infection. Using human induced pluripotent stem cells as a clinically relevant model, we show that metabolic reprogramming persists even in the absence of overt cytopathic effects and can reveal functional pathways that regulate viral replication. Our findings highlight host metabolite profiling as a complementary diagnostic strategy that may improve detection of flavivirus exposure, particularly in asymptomatic individuals or settings where conventional molecular and serologic tests are insufficient.

## Introduction

The Flavivirus genus consists of >70 virus species, many of which are primarily transmitted to human by mosquitoes and ticks (1). Among these, mosquito-borne flaviviruses such as Dengue (DENV), Zika (ZIKV), Yellow Fever (YFV), West Nile (WNV), and Japanese encephalitis (JEV) pose significant public health threats (2). West Nile virus (WNV) remains the most clinically relevant mosquito-borne virus in the continental United States, with 1,791 cases reported, including a high proportion of neuroinvasive disease and 164 deaths in 2024 (https://www.cdc.gov/west-nile-virus/data-maps/historic-data.html (3)). Dengue virus (DENV) cases are also rapidly increasing. In 2024, more than 13 million cases of dengue have been reported in North, Central, and South America and the Caribbean including 10,291 dengue cases in the U.S., 6,550 of which were locally acquired in USA (https://www.cdc.gov/dengue/data-research/facts-stats/historic-data.html (4), https://www3.paho.org/data/index.php/en/mnu-topics/indicadores-dengue-en.html (5)). Conversely, while Zika virus (ZIKV) activity has been decreasing since the 2016–2017 epidemic, its geographic footprint has been spreading. There were 224,137 ZIKV cases reported to Pan American Health Organization (PAHO) between 2019 and 2025, including over 44,445 cases occurring in 2024 in 52 countries and territories of the Americas (2) (https://www.paho.org/en/arbo-portal/zika-data-and-analysis (6)). These trends underscore the ongoing risk posed by flaviviruses in the U.S., particularly in the context of climate change, global travel, and vector range expansion (7–9).

Although flaviviruses are primarily transmitted to humans by arthropod vectors such as mosquitoes and ticks, several non-vector-borne routes of transmission have also been documented. These include sexual transmission, vertical transmission from mother to fetus, and the use of contaminated human donor tissues (10–12) Transmission through non-vector is particularly problematic as approximately 80% of individuals infected with DENV, WNV, or ZIKV remain asymptomatic, complicating surveillance, diagnosis, and threatening the safety of human tissues based therapies (2, 13).Current diagnostic strategies for flavivirus infections largely rely on nucleic acid amplification tests (NAT) and serological assays. However, both approaches face limitations. NAT is constrained by the transient nature of viremia and the often-low levels of viral RNA in clinical samples, particularly in asymptomatic or late-phase infections (14, 15). Further, serological testing is complicated by the high degree of antigenic similarity among flaviviruses, which leads to antibody cross-reactivity and reduces the specificity of differential diagnosis, a critical factor for ensuring appropriate treatment of infected individuals (16, 17). These diagnostic challenges warrant a need for more sensitive, specific, and durable test methods to detect flavivirus infections.

Metabolomics, the comprehensive study of small molecules, metabolites, in biological systems, offers a powerful window into the biochemical alterations induced by viral infection (18, 19). Flavivirus infection significantly perturbs host metabolic pathways including lipid metabolism, nucleotide biosynthesis, and energy production creating unique and detectable metabolomic fingerprints (20–22). Because these changes occur early in infection, can be detected even in the absence of overt symptoms, and can persist, making metabolomics particularly suitable for identifying asymptomatic carriers, a critical gap in current surveillance efforts. Moreover, metabolomic signatures have been identified in readily accessible biospecimens such as serum, plasma, and urine, facilitating non-invasive diagnostics (23, 24).

In this study, we leveraged metabolomics as a tool to differentiate between DENV and ZIKV flaviviruses in human induced pluripotent stem cells (hiPSCs). hiPSCs were chosen because they are widely used to model human development and diseases, perform drug screening, and develop cell therapies (25). Differentially identified metabolites identified in this study can be extrapolated to develop biomarkers for detecting flaviviruses in donor tissues. We show that DENV and ZIKV infections induce distinct, virus-specific metabolic reprogramming in hiPSCs during both acute (within 96 hours post-infection) and long-term phases (75 days post-infection). DENV3 triggered early, sustained metabolic activation, while ZIKV-MR766 caused transient suppression, reflecting divergent host responses. Long-term infections with DENV2, DENV3, and ZIKV-PRV produced unique yet overlapping metabolic signatures, with consistent alterations in amino acids, lipids, and nucleosides. Tryptophan metabolism emerged as one of the key regulatory pathways, where inhibition of the kynurenine and serotonin branches differentially impacted viral replication. These findings position metabolomics as a promising tool for flavivirus detection and for uncovering potential host-virus interactions.

## Materials and Methods

### Cell culture

Human induced pluripotent stem cell line (CL1) was generously provided by Dr. Zhaohui Ye at CBER, FDA, Silver Spring, MD. This cell line was formerly generated from peripheral blood CD34+ cells using episomal plasmid transfection (26, 27). CL1 cell line was maintained in six well plates coated with Matrigel (Cat# 354277, Corning, USA) in mTeSR™ Plus medium (Cat # 100-0276, STEMCELL Technologies). Vero cells (ATCC CCL-81) were maintained in RPMI 1640 media supplemented with 10% fetal bovine serum (Cat# A15-201, PAA Laboratories, USA), 1% penicillin-streptomycin (Cat# 30-002-CI, Corning, USA). Both cell lines were maintained at 37°C and 5% CO_2._

### Viruses

ZIKV strains (MR766, PRV) and DENV strains (DENV2, DENV3) from BEI Resources were propagated in Vero cells as described previously (28). Briefly, T75 flasks with 90% confluent Vero cells in RPMI 1640 + 2% FBS were infected at MOI 0.01-0.1 and incubated at 37°C. When 80% of cells showed cytopathic effects, supernatants were harvested by centrifugation, filtered, aliquoted, titered, and stored at −80°C as working stocks for acute and LTI experiments.

### Infections

hiPSC was maintained on Matrigel coated six well plates in mTeSR™ medium and passaged with Versene EDTA (Cat # 15040066, Gibco) every 4-5 days for 25 passages. For acute infection study, cells at 50-70% confluency were dissociated with Accutase (Cat #7920 STEMCELL Technologies, USA) and seeded at 75,000-100,000 cells/well in Matrigel-coated 6-well plates with mTeSR + 10 μM ROCK inhibitor (Cat# Y-27632, STEMCELL Technologies, USA). The medium was replaced in 24 hours to remove the ROCK inhibitor, and cells were grown for another 24 hours before viral infection at MOI 1. For infections, ZIKV MR766 and DENV 3, stocks were diluted to the MOI 1 in RPMI containing 2% FBS and added to cells. After the viral infection, medium was changed after every 48 hours. Supernatants and pellet samples from control (uninfected) and infected cells were collected at 0-, 8-, 24-, 48-, 72- and 96- hpi during acute infection. To study inhibition of tryptophan catabolism to the kynurenine and serotonin, the acute viral infection with DENV3 or ZIKV-MR766 was performed at MOI 1 for 2 hours followed by treatment with 50 µM of either 1-methyl tryptophan (1-MT) an IDO inhibitor (Cat# 452483, Sigma Aldrich) or 4-Chloro-DL-phenylalanine (4-CDP) tryptophan hydroxylase 1 and 2 inhibitor (Cat# C6506, Sigma Aldrich, USA). After 48-and 96- hours of incubation, cell pellets were collected for IFA and qRT-PCR with three biological replicates. For LTI, cells were infected with DENV2, DENV3, ZIKV-MR766, and ZIKV-PRV at MOI 1 and maintained by passaging every 4-5 days for 75 days, with infectivity confirmed by immunofluorescence assay every 4-5 passages. Infections were performed in triplicate in six well plates. Cell viability was assessed using Countess 3 (Thermo Fisher, USA), and morphology/CPE were monitored daily. Samples for metabolomics, RNA, and protein analysis were stored at −80°C for further analyses according to study overview (Fig. 1A).

**Figure 1:**
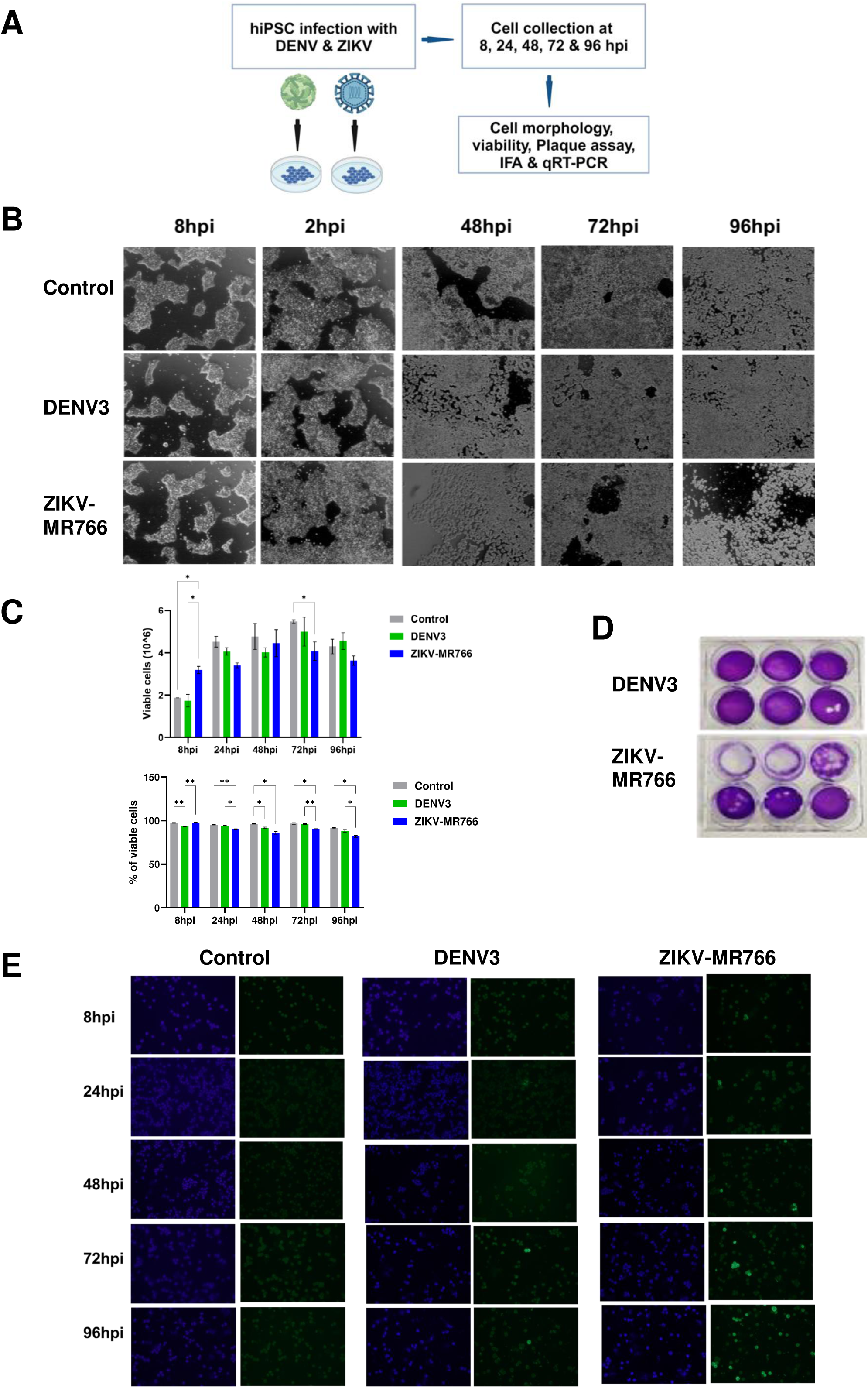
hiPSCs show differential susceptibility to DENV3 and ZIKV-MR766 infections. (A) Study overview-hiPSCs were infected with DENV3 and ZIKV-MR766 strains at MOI 1. Samples were collected 8-, 24-, 48-, 72- and 96-hours post infection (hpi). (B) Changes in morphology of hiPSCs (Control, DENV3 and ZIKV-MR766) were captured using an inverted phase contrast microscope-100x. (C) Number and percent of viable cells were determined by trypan blue dye. Significantly different samples at each time point are indicated with asterisks (p<0.05). (D) Plaque assay to study infectious virions released from hiPSCs at 96 hpi. (E) Immunofluorescence assay (IFA) with anti-4G2 antibody (green). DNA was stained with DAPI.

### Immunofluorescence assay (IFA)

To prepare cell samples for IFA, cell pellets were washed with PBS, resuspended in 500 μl PBS, and 10-20 μl was spotted onto glass slides. Slides were air-dried and fixed with methanol:acetone (1:1) for 15 minutes at room temperature. Anti-flavivirus group monoclonal antibody (Clone D1-4G2-4-15) was used as primary antibody to detect the presence of ZIKV and DENV antigens. Cells were blocked for 30 minutes with blocking solution (KPL,, USA) and then incubated with primary antibody (1:400 in 1% BSA-PBS) for 1-hour at room temperature. After PBS washing, cells were incubated with Alexa Fluor 488-conjugated goat-anti-mouse IgG secondary antibody (1:1000) for 30 minutes, washed again, and mounted with glycerol-PBS containing DAPI. Images were captured using an Olympus BX51 immunofluorescent microscope.

### Western blot

Western blot analyses followed protocol from (29). Cell pellets from acute and LTI were lysed in RIPA buffer (Cat# 89901, Thermo Scientific, USA) with protease inhibitors (Cat# 11 836 153 001, Roche) on ice for 1 hour, then sonicated at 40% power for 3 minutes. Protein concentrations were determined by bicinchoninic acid assay (BCA) (Cat# 23225, Thermo Fisher, USA) in duplicate. Samples were normalized by total protein concentration for respective time point groupings prior to loading. Normalized samples were treated with 5% 2-mercaptoethanol in Laemmli buffer, heated at 95°C for 5 minutes, and run on 4-15% SDS gels (Cat# 4568085, Bio-Rad) at 160V for 50-55 minutes. Proteins were transferred to PVDF membranes using iBlot2 (25V, 7 minutes), then blocked in 5% nonfat milk in TBS-T. Membranes were incubated with primary antibodies, anti-beta actin (Cat# 026-42210, LI-COR, USA), anti-Dengue NS1 (Cat# MAB94441-100, R&D Systems, USA) and anti-Zika envelope (Cat# GTX133314, GeneTex, USA) for 1 hour at room temperature, washed 3x with TBS-T, incubated with peroxidase-labeled secondary antibodies (Cat# 5450-0010, #214-1806 KPL, USA) for 30 minutes, and washed again. Detection used Pico or Femto developing reagents on an Amersham Imager 600.

### Plaque Assay

Vero cells were cultured in 6-well tissue culture plates and allowed to reach 80-90% confluency to form complete monolayers. The confluent cell monolayers were then infected with serial 10-fold dilutions of viral supernatants for 2 hours at 37°C to allow for optimal viral attachment and entry. Following the inoculum was replaced with an agar overlay medium and plates were incubated at 37°C in an incubator for 5 days to allow sufficient time for viral replication and plaque development. Visible plaques became apparent on the 5th day post-infection, at which point the plates were stained with crystal violet solution for plaque visualization (30).

### qRT-PCR

Quantification of viral RNA from DENV3 and ZIKV-MR766 infected samples, was achieved by real-time reverse transcription polymerase chain reaction (RT-PCR). Total RNA was extracted from pellets of control and infected samples using PureLink RNA Mini kit (Cat# 12183025, Invitrogen) followed by quantification utilizing a Qubit 4 fluorometer with the RNA High Sensitivity kit. ZIKV RNA quantification was conducted using the iTaq Universal SYBR Green One-Step Kit (Bio-Rad, USA), with each 20 μL reaction mixture containing 3-5 μL of extracted RNA template, 10 μL of iTaq universal SYBR Green reaction mix, 0.25 μL of iScript reverse transcriptase, 1 μL each of 5 μM forward (CCTTGGATTCTTGAACGAGGA) and reverse (AGAGCTTCATTCTCCAGATCAA) ZIKV-specific primers previously published by (31) and (28).

DENV3 detection was accomplished using a TaqMan-based real-time RT-PCR assay following the protocol established by (32) utilizing primer sequences DEN-3 F (GGACTGGACACACGCACTCA), DEN-3 C (CATGTCTCTACCTTCTCGACTTGTCT), and the corresponding TaqMan probe (ACCTGGATGTCGGCTGAAGGAGCTTG) for specific viral genome detection.

### Sample preparation for metabolomics (LC-MS)

Chemicals: Water, methanol, and 2-propanol, both CHROMASOLV LC-MS grade, buffer additives for online mass referencing, media, and sample preparation chemicals at the highest available purity were purchased from Sigma-Aldrich and Agilent Technologies. Pure water for extraction and resuspension was obtained with an electric resistance of greater than 16 MΩ from a NANO pure purification unit (Barnstead, Dubuque, United States). Standard solutions were prepared as described previously by Büscher et al., (33).

### Acute Infection LC-MS

Untargeted metabolite analysis was conducted on a QExactive HF-X mass spectrometer equipped with a HESI II probe. The mass spectrometer was coupled to a Vanquish binary UPLC system (Thermo Fisher Scientific,, USA). For chromatographic separation prior to mass analysis, 5 µL of the sample was injected onto a BEH Z-HILIC column (100 mm, 1.7uM particle size, 2.1 mm internal diameter, Waters). Samples were diluted 1:5 in acetonitrile for the analysis. Each sample was analyzed twice in two different chromatographic separations. In method 1, Mobile phase A was 15 mM ammonium bicarbonate in 90% water and 10% acetonitrile, and mobile phase B was 15mM ammonium bicarbonate in 95% acetonitrile and 5% water. In Method 2, Mobile phase A consisted of 10 mM ammonium formate in 50/50 Acetonitrile/Water, brought to pH 2.7 with formic acid, and Mobile phase B was 10 mM ammonium formate in 95/5/5 Acetonitrile/Methanol/Water at pH 2.7. Mass analysis was conducted in positive mode for Method 2 and negative mode for Method 1.

For both methods, the column oven was held at 45°C and autosampler at 4°C. The chromatographic gradient was carried out at a flow rate of 0.5 ml/min as follows: 0.75 min initial hold at 95% B; 0.75-3.00 min linear gradient from 95% to 30% B, 1.00 min isocratic hold at 30% B. B was brought back to 95% over 0.50 minutes, after which the column was re-equilibrated under initial conditions. The mass spectrometer was operated in full-scan negative mode, with the spray voltage set to 3 kV (3.5 kV for positive mode), the capillary temperature to 320 °C, and the HESI probe to 300 °C. The sheath gas flow was set to 50 units, the auxiliary gas flow was set to 10 units, and the sweep gas flow was set to 1 unit. Mass acquisition was performed in a range of m/z = 70–900, with the resolution set at 120,0000. Retention times were determined using authentic standards. Raw data were converted to mzML files using msConvert ver.3.0.22046-caef294 (34).

### Data Processing

For positive ionization mode, 85 mzML files associated with blank, NIST, quality control sample which is a pool of all study samples (pSS), and sample injection were uploaded into MZmine v3.0.11(35) for peak extraction and default parameters were used. A total of 10,077 features were extracted by MZmine. Features were filtered on 3 parameters: missing data, signal-to-noise ratio (S/N), and coefficient of variance (CV). Features with missing values in >20% of samples were removed. S/N ratios were calculated between the QC injections and blank injections and any feature with a S/N values <3 was removed. Additionally, any feature with a CV >20% across all QC samples were removed. After filtration, 1,636 features remained.

For negative ionization mode, 78 mzML files associated with blank, NIST, and sample injection were uploaded into MZmine v3.0.11 (35) to peak extraction and default parameters were used. A total of 7,076 features were extracted by MZmine. Features were filtered on 2 parameters, missing data and S/N using the same criteria as for the positive ionization mode data (e.g., removal of features with missing values in >20% of samples and S/N values <3). After filtration, 2,254 features remained for further analysis.

Data were normalized to biomass, missing values were imputed with minimum intensity value for a given metabolite across all samples divided by 2, and intensities were log10 transformed. Data quality via PCA plots showed good clustering if blank, NIST, and pSS injection indicating good quality of data (Fig. S2A-B). Identities of metabolites were determined by comparing fragmentation spectra to online available databases such as - (36) and LIPID MAPS-(37).

### Persistent Infection LC-MS

#### Reagents and solvents

LC-MS grade water, acetonitrile (ACN), isopropanol (ISP), methanol (MeOH), acetic acid (AcOH, > 98% pure), formic acid (>99% pure) were purchased from ThermoFisher Scientific (USA). LC-MS grade Ammonium formate and ammonium acetate were purchased from Fluka Honeywell (Muskegon, MI). UHPLC-grade methyl tert-butyl ether (MTBE) was purchased from Fisher Scientific (, USA). Mass Spectrometry Metabolite Library of Standards (MSMLS) was purchased from IROA Technologies (USA).

#### Lipid concentration from cell extract

Cell extracts were obtained from 1 million cells, using 1 mL of a solvent mixture composed of 40% methanol (MeOH), 40% acetonitrile (ACN), and 20% water. To concentrate the lipids, a biphasic extraction was performed on the cell extracts using methyl tert-butyl ether (MTBE) and MeOH, following a protocol adapted from a previously published method.

Briefly, 50 µL of the cell extract was mixed with 30 µL of water and 160 µL of MTBE. The samples were vortexed for 2.5 minutes, followed by centrifugation for 2 min (10,000g, 4°C). The supernatant was carefully collected and dried under a stream of nitrogen gas. This extraction protocol was performed twice to ensure optimal recovery of lipids.

#### UHPLC-MS Analysis for Metabolomics and Lipidomics

Lipid and polar metabolite extracts were analyzed using an Agilent 1290 Ultra High-Performance Liquid Chromatograph (UHPLC) coupled to an Agilent 6546 Quadrupole Time-of-Flight Mass Spectrometer (Q-TOF MS) (Agilent Technologies,, USA). For lipidomics analysis, lipid extracts were reconstituted in a solvent mixture of acetonitrile/isopropanol (ACN/IPA, 50 µL, 7:3, v/v) and vortexed for 2.5 minutes. For metabolomics analysis, 50 µL of polar extracts were diluted in 50 µL of ACN to reach a 90% organic phase to ensure optimal peak shape before analysis.

Lipid separation was performed using a previously published method (38)on a C8 column (Acquity Plus BEH, Waters, Milford, MA, 100 mm × 2.1 mm, 1.7 μm particle size) with an injection volume of 3 µL. Polar compound separation was performed using a modified protocol based on (39), with solvent modifications according to (40), on a HILIC column (Atlantis, Waters, Milford, MA, 150 mm × 2.1 mm, 3 µm particle size) with an injection volume of 5 µL.

For both lipidomics and metabolomics, the mobile phases used were phase A (95% water, 5% ACN, 0.1% acetic acid, 0.1% ammonium acetate for lipids and 0.2% formic acid, 20 mM ammonium formate for polar compounds) and phase B (95% ACN, 5% water, 0.1% acetic acid, 0.1% ammonium acetate for lipids and 0.2% formic acid for polar compounds). The gradient was run at 0.4 mL/min for lipid separation and 0.25 mL/min for polar metabolite separation. For lipidomics, the chromatographic gradient included the following: 55% B held for 1 min, followed by a linear increase to 65% B over 3 min, a linear increase to 85% B over 8 min, a linear increase to 99% B over 3 min, holding at 99% B for 3 min, and then rapid re-equilibration to initial conditions until 22 min. For metabolomics, the gradient included a 5% A held for 1 min, followed by a linear increase to 40% A over 6.4 min, a linear increase to 60% A over 0.1 min, holding at 60% A for 5 min, and then rapid re-equilibration to the initial conditions until 22.7 min. To reduce the equilibration time of the HILIC column, the flow was increased to 0.5mL/min between 14.6 and 21.6 min.

Source parameters for both positive and negative ion modes were as follows: VCap = 3000 V, gas temperature = 300°C, drying gas flow rate = 10 L/min, nebulizer pressure = 25 psi, sheath gas temperature = 350°C, and sheath gas flow rate = 11 L/min. The mass spectrometer was tuned and calibrated before analysis and throughout each sample injection, using the manufacturer’s reference mixture for calibration.

Pooled quality control (QC) samples, consisting of equal aliquots from each sample within the same extraction type either polar or apolar were analyzed every 5 samples to monitor instrument performance, and solvent blanks were analyzed at the beginning of the sequence, before samples or QC, to remove any contaminant features from the data.

The identity of significant compounds was confirmed by comparing their mass to charge ratio (m/z) and retention times (RT) with purified standards to achieve a Level 1 identification, or by fragmenting the compounds using three different collision energies (10, 15, and 20 eV) and matching the resulting fragments with online spectral libraries for Level 2 identification, according to the guidelines set by the Metabolomics Standards Initiative (MSI (41))

#### Data Processing

Raw data were converted to mzData files using Agilent MassHunter Qualitative Analysis software (version 10.0). mzData files associated with blank, process blank, QC, and sample injection were uploaded into MZmine v2.26 (42) to peak extraction and full set of parameters can be found in Supplementary Table 1. A total of 5,823 and 903 features were extracted by MZmine for positive ionization mode and negative ionization modes, respectively. Similar to the acute infection, features were filtered on 3 parameters, missing data, S/N, and CV. Any features with missing values for >20% of samples were removed. S/N ratios were calculated between the QC injections and process blank injections and any feature with a S/N values <3 was removed. Additionally, any feature with a CV >20% in the QC samples were removed. After filtration, 1,111 and 119 features remained for positive and negative ionization modes, respectively. Data were normalized to biomass, missing values were imputed with minimum intensity value divided by 2, and intensities were log10 transformed.

The lipidomics workflow follows the same steps as the metabolomics workflow in the previous paragraph. A total of 42,065 and 2,936 features were extracted by MZmine v2.26 (42) for positive and negative ionization modes, respectively. After filtration, 1,170 and 282 features remained for positive and negative ionization modes, respectively. Data were normalized to biomass, missing values were imputed with minimum intensity value divided by 2, and intensities were log10 transformed.

### Statistical Analyses

#### Acute Infection

All statistical tests and figures were generated in R version 4.4.3 (43) T-test comparisons were performed between control and infection at each time point. For example, control vs DENV3 at 8 hpi, control vs ZIKV-MR766 at 8 hpi, control vs DENV3 at 24 hpi, control vs ZIKV-MR766 at 24 hpi, etc. P-values were adjusted for multiple comparisons using Benjamini-Hochberg (FDR). Additionally, fold changes were calculated between infection and control. Features with FDR corrected p-value < 0.3 and fold changes values >1.25 or <0.75 were considered as statistically significant.

#### Persistent Infection

Statistical tests were done in the same format as the acute infection. T-test comparisons were performed between control and infection, P-values were adjusted for multiple comparisons using FDR and fold changes were calculated between infection and control. Features with FDR corrected p-value < 0.05 and fold changes values >1.25 or <0.75 were analyzed further.

## Results

### Differential Susceptibility of hiPSCs to DENV3 and ZIKV-MR766 During Acute Infection

Undifferentiated pluripotent stem cells are known to exhibit resistance to viral infection, while differentiated derivatives are generally more permissive (44), Infection of CL1 hiPSC line (produced from CD34+ peripheral blood cells via episomal reprogramming) with DENV-3 and ZIKV showed no major morphological changes or cytopathic effects (CPE) at any tested time points as compared to negative control as shown in Fig 1B. However, cellular viability evaluation revealed higher viability at 8 hpi for those infected with ZIKV-MR766 as compared to those infected with DENV3 and non-infected control. By 72 hpi, this trend had reversed, and ZIKV-MR766 infected showed significant reduction in viability as compared to DENV-3 and non-infected control (Fig. 1C). ZIKV-MR766 infection significantly reduced the percentage (%) of viable cells as compared to both non-infected control and DENV3 at all time points except 8hpi. (Fig. 1C).

Productive infection, assessed by PFU in culture supernatants, showed robust infection of CL1 with ZIKV-MR766 demonstrated by high titer of infectious virions at 96hpi (TCID_50_ 4.41×10^7^ serially diluted 10-fold in six wells). In contrast, DENV3 resulted in non-productive infection as no infectious virions were detected by PFU (Fig. 1D). Protein production were evaluated by IFA and western blot. IFA was performed using anti-flavivirus envelope antibody (4G2). ZIKV-MR766 was detected as early as 48 hpi, with increased detection at 72 and 96 hpi, while DENV3-strain was weakly detectable at 72 and 96 hpi (Fig. 1E).

Western blotting using anti-ZIKV envelope, anti-DENV NS1, and anti-actin antibodies corroborated the IFA results. ZIKV protein was detectable by 48 hpi, while DENV3 protein was weakly detectable at 96 hpi (Fig. S1A). However, qRT-PCR revealed ZIKV RNA as early as 8 hpi, and DENV3 RNA at 48 hpi (Fig. S1B). Together, these findings confirm that undifferentiated hiPSCs are more susceptible to ZIKV-MR766 than to DENV3 during acute infection.

### Establishment and Characterization of Long-Term Flavivirus Infection in hiPSCs

To investigate long-term host-flavivirus interactions through a metabolomics lens, we established long-term infections (LTI) of DENV2, DENV3, and ZIKV-PRV in hiPSC cultures (see materials and methods). ZIKV-MR766 infected hiPSC cells failed to survive beyond the second passage, this may be attributed to ZIKV-MR766induced differentiation of hiPSCs, ultimately leading to cell death (45). There were no significant differences in morphology or cell death observed between long-term infected and uninfected control cells (Fig. 2A). Cellular viability evaluation revealed that at 48 hours, the number of viable cells was comparable across all groups. Interestingly, by 72 hours, DENV2 and DENV3 infected cells, but not ZIKV-PRV infected cells, exhibited a highly significant reduction in number of viable cells relative to control. At 96 hours, all infected groups showed a marked decrease in viable cell numbers compared to the control (Fig. 2B). Notably, percent-viability of long-term infected cells remained close to 100% at 72h and 96h, indicating that the decrease in viable cell counts reflects impaired proliferation rather than increased cell death in infected cultures. (Fig. 2B). At 48h the only DENV3 infected cells showed significantly low % of viable ells as compared to control. IFA using 4G2 confirmed high levels of infection across all long-term infected lines DENV2, DENV3 and ZIKV-PRV infected cells, demonstrating comparable antigen positivity, suggesting robust and sustained viral presence (Fig. 2C).

**Figure 2:**
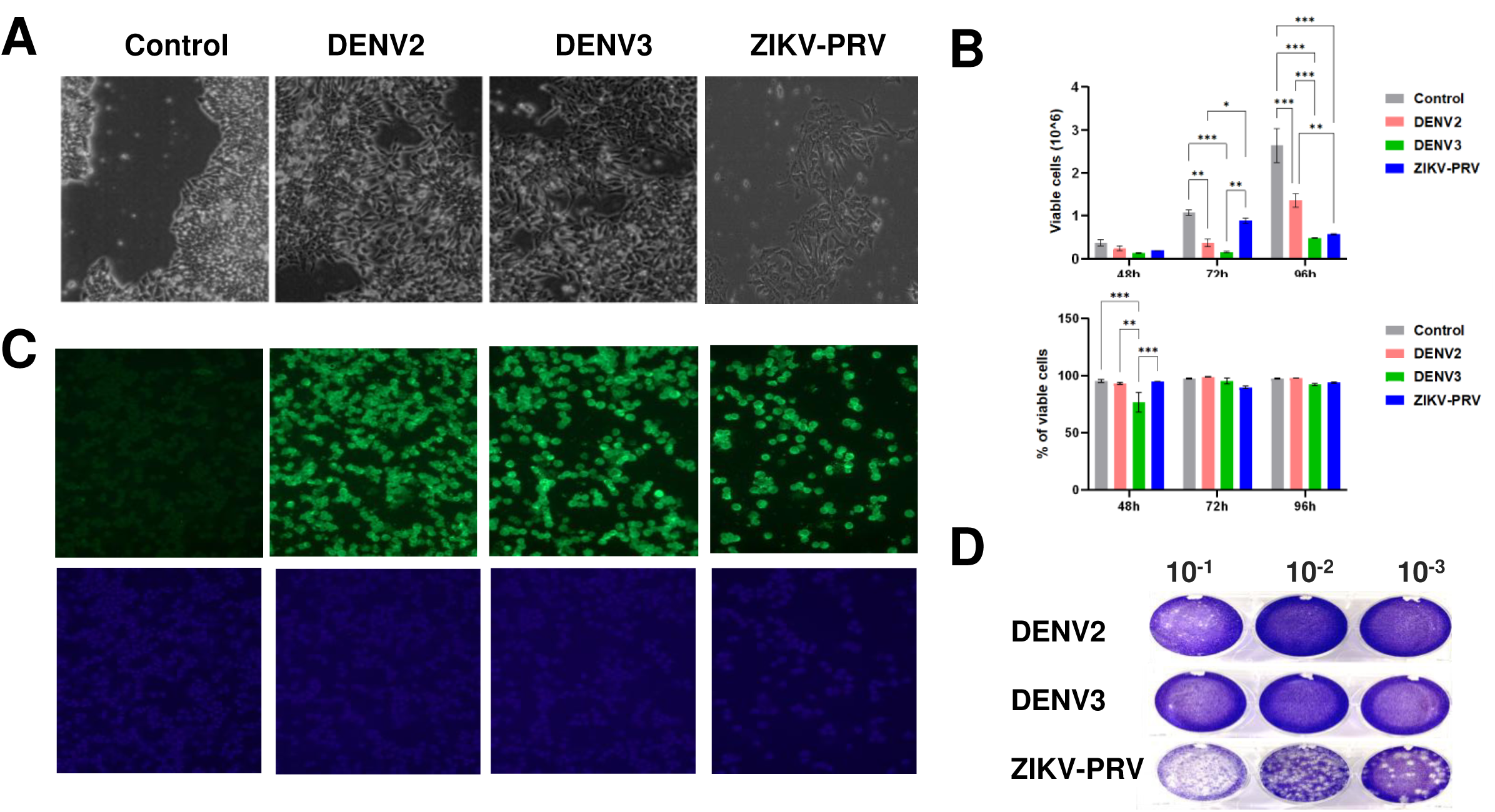
Establishment of long-term DENV2, DENV3 and ZIKV-PRV Infection in hiPSCs. Cells were passaged for 75 days post infection (dpi) and assessed at 72 hours (A) Morphology of control and LTI hiPSCs. (B) Number and percent of viable cells of viable cells were determined by trypan blue dye. Significantly different samples at each time point are indicated with asterisks (p<0.05). (C) IFA with anti-4G2 antibody (green), DNA was stained with DAPI. (D) Plaque assay to study infectious virions released from hiPSCs.

Characterization of supernatants from LTI cultures revealed virus-specific differences in productive infection. ZIKV-PRV consistently produced infectious virions across all dilutions, mirroring results from acute infection assays. In contrast, DENV2 produced a low number of infectious virions, detectable only at lower dilutions, and no plaques were observed for DENV3 (Fig. 2D) (46). These findings confirm that hiPSCs can support long-term flavivirus infection, with susceptibility varying by viral strains.

### Metabolomics differentiates DENV and ZIKV infected cells from non-infected controls upon acute infection

hiPSCs infected with DENV3 or ZIKV-MR766 were harvested at different timepoints for untargeted metabolomics analysis as outlined in Fig. 3A (see the Materials and Methods). Principal component analysis (PCA) revealed time-specific clusters and within each timepoint, infected and control samples were slightly divergent (Fig. 3B). Notably, ZIKV-MR766-infected samples at T8 appeared metabolically distinct from all other groups, suggesting an early temporal metabolic shift due to infection (Fig. 3B).

**Figure 3:**
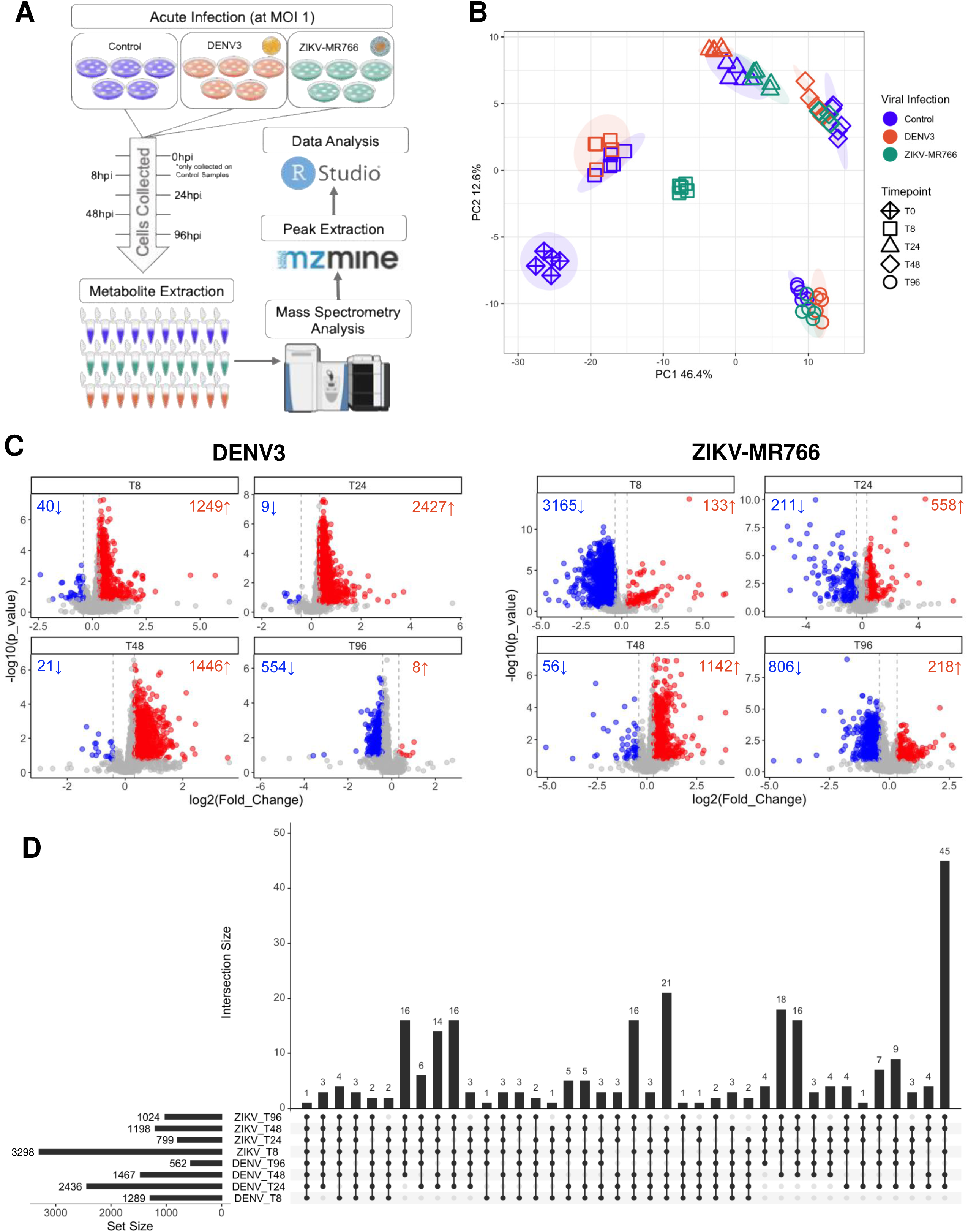
Overview of untargeted mass spectrometry-based metabolomics results for acute flavivirus infection. (A) Schematic overview of the metabolomics workflow in acute infection. (B) PCA representation of the data primarily shows separation of samples based on timepoint. (C) Volcano plots show significantly up- (red) and down- (blue) regulated features based on FDR values <0.3 and fold changes values >1.25 or <0.75. Plots are separated by viral infection and further separated by timepoint. (D) UpSet plot shows overlap of features across both viral infections and all four timepoints.

Volcano plots in Fig. 3C highlighted the temporal dynamics and virus-specific metabolic responses of DENV3 and ZIKV-MR766-infected hiPSCs (adjusted p-value<0.3 and fold change (FC) >1.25 or <0.75). DENV3-infected cells showed a predominant upregulation of features at early timepoints, with 1,249, 2,427, and 1,446 features significantly upregulated at 8-, 24-, and 48- hpi, respectively. Downregulated features were fewer, indicating a primarily activated metabolic response. By 96 hpi, the number of significantly dysregulated features diminished, suggesting partial metabolic recovery or adaptation. In contrast, ZIKV-MR766 infection resulted in a striking early downregulation of features, with 3,165 features significantly downregulated and only 133 upregulated at 8 hpi. By 24 and 48 hpi, the number of upregulated features increased to 558 and 1142, while 56 and 218 features remained significantly downregulated, indicating a shift toward a more balanced or compensatory metabolic state. Interestingly, at 96 hpi, the pattern was similar to 8 hpi, with 806 downregulated and 218 features upregulated. These findings suggest a rapid and sustained host metabolic reprogramming in response to ZIKV-MR766, distinct from the DENV3 response in hiPSCs.

An UpSet plot showing the top 40 sets of significantly altered features across all eight infection/timepoint comparisons shows that 145 features were consistently altered in six or more comparisons (Fig. 3D), underscoring shared host responses across viruses and timepoints. UpSet plots for viral specific features show some overlap across time points with a majority of features are time-point specific or significant across two time points (Fig. S2C-D). Notably, suberic acid was the only metabolite differentially regulated across all eight comparisons. Further, metabolite annotation revealed 23 metabolites differentially regulated in five or more conditions, including 12 amino acids, peptides, and analogues (e.g., glutamic acid, arginine, tryptophan), 4 lipids (e.g., suberic acid, glycerophosphoserine), 3 nucleosides, and 4 other small molecules (Fig. S3).

### Metabolomics and lipidomics differentiates hiPSC post Long-Term infection (LTI) with DENV2, DENV3 and ZIKV-PRV

Polar and lipid fractions from LTI samples were run in both ionization modes. In the polar fraction, LC-MS yielded 13,697 and 489 features in respective positive and negative ionization modes. In the lipid fraction, there were 42,065 and 2,938 features in positive and negative ionization modes, respectively. Unlike acute infection samples, PCA of the metabolite profiles demonstrated clear separation between infected and control samples, indicating distinct metabolic alterations induced by each virus in long term (Fig. 4B and S4A-D). ZIKV-PRV showed the most divergent profile from the control group. Notably, despite being strains of the same virus, DENV3 clustered more closely with ZIKV-PRV than with DENV2, suggesting shared long-term metabolic signatures between these two viruses (Fig. 4B and S4A-D).

**Figure 4:**
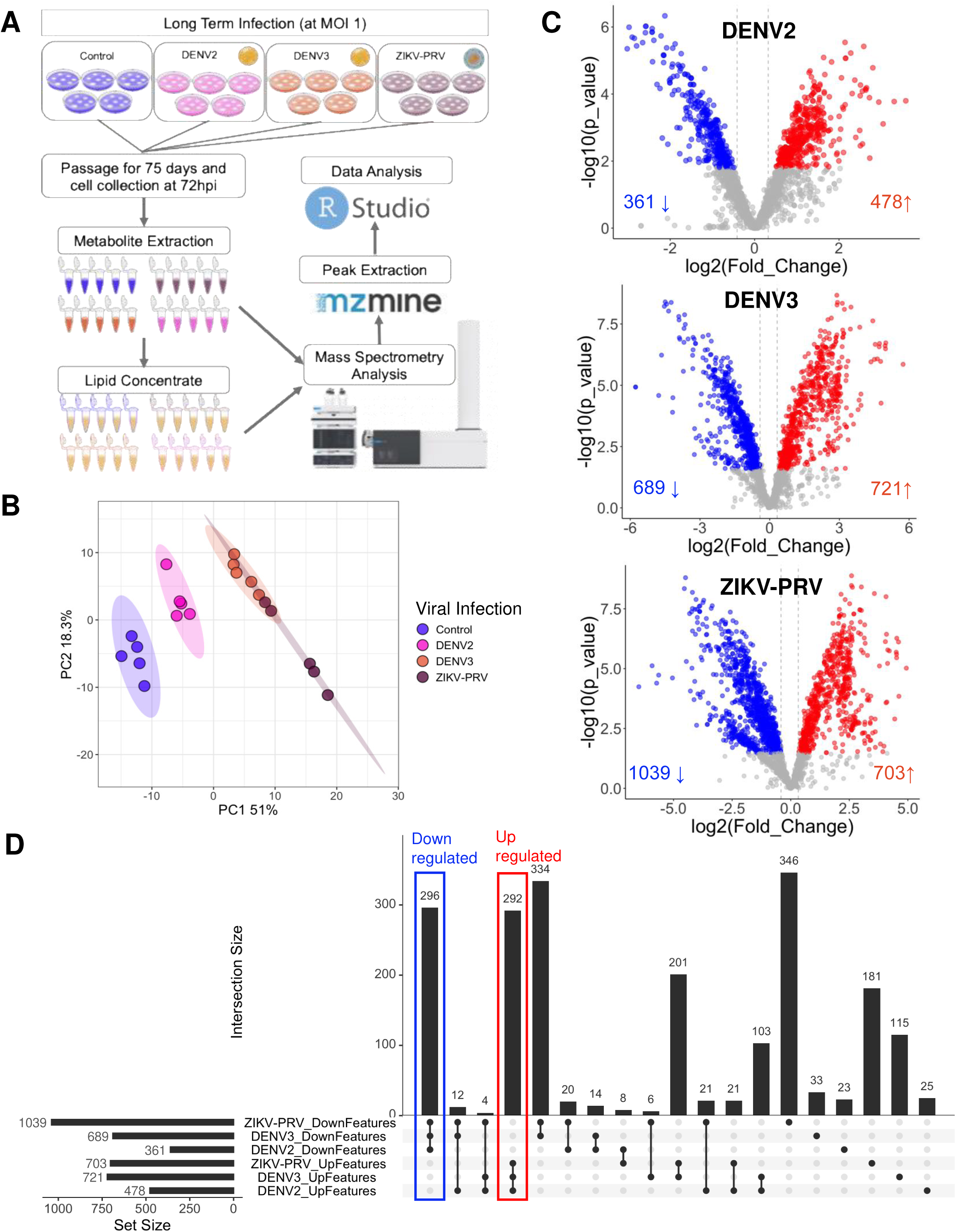
Overview of untargeted mass spectrometry-based metabolomics results for long term flavivirus infection. (A) Schematic overview of the LTI metabolomics workflow. (B) PCA representation of the data shows separation of samples based on viral infection. (C) Volcano plots show significantly up- (red) and down- (blue) regulated lipid (circle) and polar (triangle) features based on FDR values <0.05 and fold changes values >1.25 or <0.75. Plots are separated by viral infection. (D) UpSet plot shows overlap of features across all three viral infections and whether that feature was significantly up or down regulated.

Volcano plot analysis identified hundreds of differentially regulated features for each infection type using a threshold of adjusted p < 0.05 and (FC) >1.25 or <0.75 (Fig. 4C). DENV2 infection led to 478 upregulated and 361 downregulated features, while DENV3 yielded 721 upregulated and 689 downregulated features. ZIKV-PRV induced the most extensive changes, with 703 upregulated and 1,039 downregulated features (Fig. 4C). Next, we performed an UpSet analysis, to identify common and unique features, (Fig. S4E) while also taking into account the directional change of the feature (Fig. 4D). This revealed 292 features consistently upregulated and 296 consistently downregulated across all three infections. Pairwise comparisons showed greater overlap between DENV3 and ZIKV-PRV (334 downregulated; 201 upregulated features) than between DENV3 and DENV2 (14 downregulated; 103 upregulated) or DENV2 and ZIKV-PRV (20 downregulated; 21 upregulated). These results suggest that long-term infection induces conserved metabolic shifts, particularly between DENV3 and ZIKV-PRV. Identification of those features that overlapped across all three infections showed 9 amino acids, peptides, and analogues; 3 nucleosides, nucleotides, and derivatives; and 62 lipids and lipid-like molecules (Fig. S5).

Looking generally across both experiments, there is a theme of three chemical classes being altered: 1) amino acids and associated metabolites; 1) nucleosides and associated metabolites, and 3) lipid and lipid-like molecules. Of all the metabolites identified in acute infection and LTI, six metabolites were found to have significantly changed between control and infected for at least one virus: alanyl-glutamine, glutamate, N-Acetylputrescine, N-N-dimethylarginine, thiamine and tryptophan (Fig. 5A).

**Figure 5:**
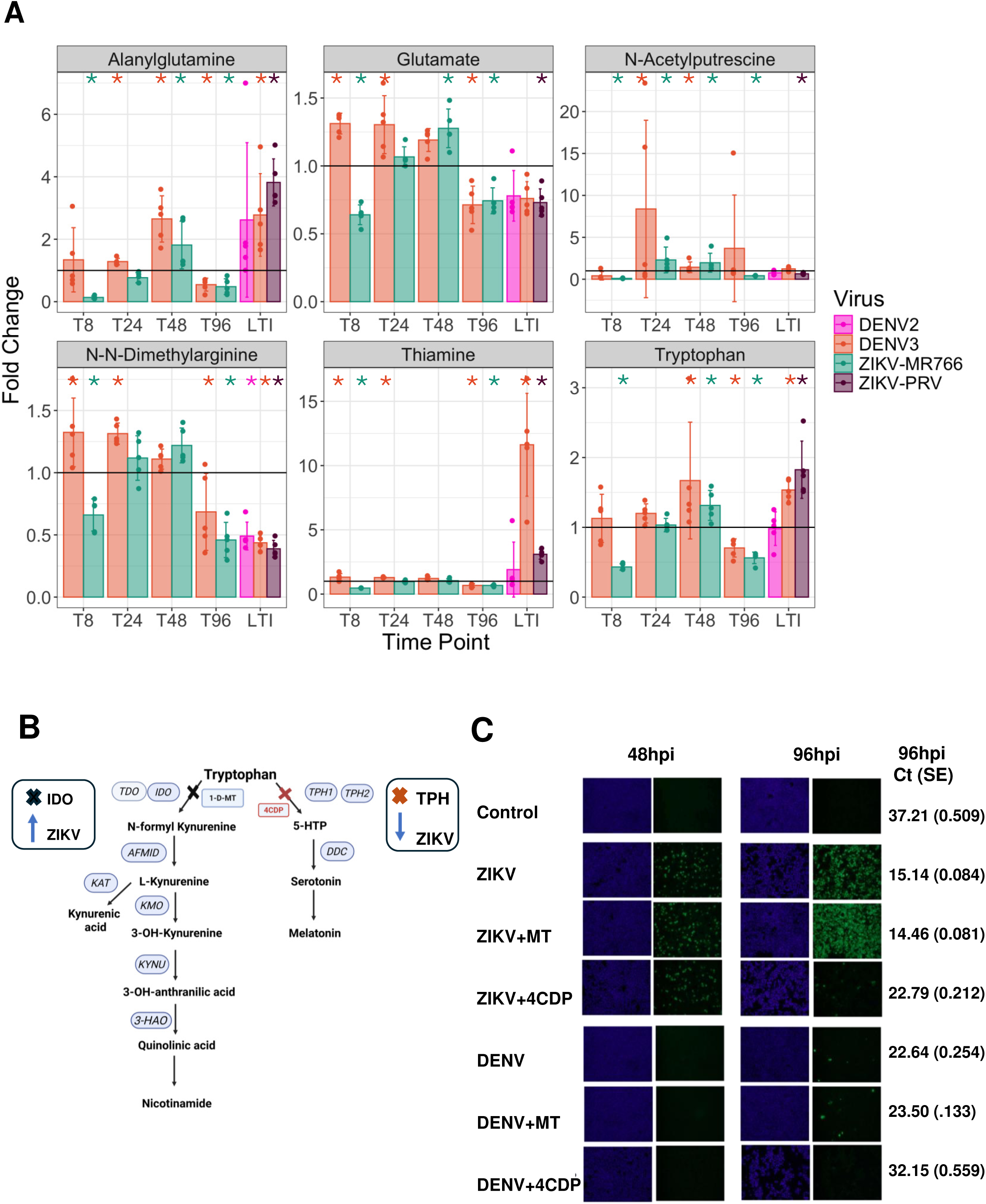
Overlap of metabolites across acute and LTI and validation studies. (A) Barplot fold-change representation of overlapping metabolites in acute and LTI. Asterisks denote those with significant p-values. (B) Schematic illustration of the tryptophan metabolic pathway highlighting the effects of pathway-specific inhibitors. 1-Methyl-D-tryptophan (1-D-MT), shown in black, inhibits the IDO enzyme, leading to increased ZIKV infection in hiPSCs. In contrast, 4-Chloro-DL-phenylalanine (4-CDP), marked in red, inhibits tryptophan hydroxylase (THP), resulting in reduced IKV and DENV load. (C) IFA with anti-4G2 after inhibition of IDO and TPH1; qRT-PCR (Ct values with standard error).

### Tryptophan catabolism regulates DENV and ZIKV infection in hiPSCs

Since tryptophan was identified as one of the significantly regulated metabolites, we next specifically evaluated two major branches of tryptophan catabolism, the kynurenine and serotonin pathways, in the context of DENV3 and ZIKV-MR766 infection in hiPSC cells (Fig. 5B). hiPSCs infected with ZIKV-MR766 or DENV3 (MOI 1) were treated with 50 µM of either 1-MT (indoleamine 2,3-dioxygenase (IDO) inhibitor) or 4-Chloro-DL-phenylalanine (tryptophan hydroxylase 1 and 2 inhibitor). After 48- and 96-hours post incubation, cells were analyzed for viral infection using IFA (Fig. 5C). Interestingly, treatment with 1-MT resulted in increased ZIKV-MR766 infectivity at 96 hpi, as evidenced by a higher number of IFA-positive cells compared to untreated controls. In contrast, inhibition of serotonin biosynthesis via 4-CDP treatment led to a marked decrease in the number of infected cells. These findings underscore the importance of tryptophan metabolism, particularly the serotonin pathway, in decreasing susceptibility to flavivirus infection (Fig. 5C). We confirmed our IFA results with qRT-PCR which showed consistent results at 96 hours post-infection (Fig. 5C, Ct values). Similar effects were observed in DENV-infected hiPSCs treated with 4-CDP, further supporting the role of serotonin pathway modulation in infection outcomes (Fig. 5C).

## Discussion

Flaviviruses remain a global public health challenge, with rising incidence, expanding geographic range, and high rates of asymptomatic infection complicating diagnosis and control (2, 13). Standard diagnostics using nucleic acid testing and serology face significant limitations due to short viremia windows and antibody cross reactivity (2, 14, 17), particularly in endemic regions with co-circulating flaviviruses. Our study helps fill this diagnostic gap by showing that untargeted metabolomics not only identifies commonly dysregulated metabolites in acute and LTI, but also reliably differentiates DENV and ZIKV infections through virus specific metabolic reprogramming in hiPSCs.

Our work highlights both acute and LTI phases, capturing dynamic and durable changes in host metabolism. During acute infection, DENV3 induced sustained metabolic activation, whereas ZIKV-MR766 caused early metabolic suppression, revealing distinct host responses. LTI models with DENV2, DENV3, and ZIKV-PRV showed virus-specific yet overlapping metabolic signatures, even in the absence of cytopathic effects, consistent with reports of subclinical flaviviral persistence in neural progenitor and other cell types (47–55). Importantly, ZIKV-PRV and DENV3 shared more metabolomic overlap than DENV2 after LTI, underscoring the role of viral strain in shaping host-pathogen interactions. A core set of metabolites including tryptophan, glutamate, and thiamine was consistently altered across infection types and timepoints. These metabolites are thus of particular interest as potential biomarkers of DENV, ZIKV and maybe other flavivirus infection.

These molecules are critical to immune regulation, neurotransmission, and cellular stress responses (19, 56, 57) and may serve as stable biosignatures for flavivirus detection even in latent or asymptomatic phases. These findings are strongly supported by previous reports that flavivirus infection perturbs central carbon metabolism, including glycolysis, the TCA cycle, and amino acid metabolism (20, 58, 59).

Tryptophan metabolism emerged as a key regulatory node as: inhibition of the discovered serotonin pathway via 4-CDP reduced the viral load. These findings align with recent evidence that serotonin supports replication of RNA viruses (60, 61).

In addition to amino acid metabolism, lipid sub classes lysophosphatidylcholine (LPC), lysophosphatidylethanolamine (LPE), phosphatidylcholine (PC), PC plasmalogen, phosphatidylethanolamine (PE), phosphatidylinositol (PI), ceramides (Cer), dihexosylceramide (Hex2Cer), sphingomyelin (SM), and cholesterol were differentially regulated in LTI samples (Fig. S5B). Metabolites involved in LPE, PE, PC, ceramide, and sphingomyelins were significantly altered in both our hiPSC model and previous ZIKV-and DENV-infected samples (62–64). While earlier research has typically focused on acute infection and single virus systems, our study uniquely integrates acute and long-term infection phases and directly compares DENV and ZIKV strains side-by-side in the same human cellular background, enabling virus- and strain-specific metabolic signatures to be resolved with high confidence.

From a public health perspective, this work provides a path forward for improving flavivirus surveillance, particularly in asymptomatic populations and tissue donor screening. As climate change, global travel, and urbanization expand arbovirus transmission zones (7, 65), the development of non-invasive, metabolomics based diagnostics and therapeutics becomes increasingly urgent. Future studies should validate these signatures in plasma or urine from infected individuals and expand to other clinically relevant flaviviruses.

## Acknowledgments

The authors thank Drs. Bharat Joshi and Alan Baer for critical reading of this manuscript and helpful comments. We thank Mr. Adam Tisch from NCATS for help with sample processing of long-term infection samples for metabolomics. This research was supported in part by the Intramural Research Program of the National Institutes of Health (NIH). The contributions of the NIH author (s) were made as part of their official duties as NIH federal employees, are in compliance with agency policy requirements, and are considered Works of the United States Government. However, the findings and conclusions presented in this paper are those of the author(s) and do not necessarily reflect the views of the NIH or the U.S. Department of Health and Human Services.

## Funding

This work was supported by the Intramural Research Program of the Center for Biologics Evaluation and Research (CBER), U.S. Food and Drug Administration. This project was also supported in part by Aaron Scholl’s appointment to the Research Participation Program at CBER administered by the Oak Ridge Institute for Science and Education through the US Department of Energy and U.S. Food and Drug Administration. This work was supported in part by the intramural program Metabolomics and multiomics (ZIC TR000547) at the National Center for Advancing Translational Sciences, part of the National Institutes of Health.

## Author Contributions

Conceptualization: TF, SD

Methodology: TF, KM, DB, AS, BL, EM, SD

Investigation: TF, KM, EM, SD

Visualization: TF, KM, EM, SD

Supervision: EM, SD

Writing- original draft: TF, KM, EM, SD

Writing- review & editing: TF, KM, DB, AS, BL, MR, EM, SD

## Competing Interests

The authors declare that they have no competing interests.

## Data availability statement

All data needed to evaluate the conclusions in the paper are present in the paper and/or the Supplementary Materials. Metabolomic data are available at MetaboLights (REQ20251229215726).

The ZIKV and DENV LTI hiPSC cell/data can be provided by CBER/FDA pending scientific review and a completed material transfer agreement. Requests for the long-term infected cells should be submitted to Sandip De.

## Supplementary Figures

**Figure S1**: Detection of DENV and ZIKA in infected hiPSCs. (A) Western blot analysis with anti-ZIKV envelope, anti-DENV NS1and anti-actin antibodies. (B) qRT-PCR (Ct values) for Actin, ZIKV-MR766 & DENV3 from control and infected samples.

**Figure S2:** (A) Acute PCA showing both ionization modes and QCs. (B) Acute Upset plot for each viral infection.

**Figure S3:** Acute infection study. Identified significant metabolite line plots. Differential regulation of potential markers of interest. LC-MS/MS analyses showed differential regulation of many features. Presented here 23 metabolites of interest between control vs DENV3 and control vs ZIKV-MR766 (statistical significance was calculated using t-test with an FDR correction)

**Figure S4:** Long-term infection. (A-D) PCA for Polar/Lipid showing both ionization modes and QCs. (E) UpSet plot for LTI cells lines.

**Figure S5:** Persistent identified significant metabolite/lipids as box plots (A) or FC dot plots (B).

## Supplementary Tables

**Table S1:** MZmine v2.26 Extraction Parameters for Persistent Data.

**Table S2:** Acute identified metabolites with identifying parameters (m/z, RT, mode, etc) and statistics values.

**Table S3:** Persistent identified metabolites with identifying parameters (m/z, RT, mode, etc) and statistics values.

